# Arrival and proliferation of the invasive seaweed *Rugulopteryx okamurae* in NE Atlantic islands

**DOI:** 10.1101/2021.06.25.448933

**Authors:** João Faria, Afonso CL Prestes, Ignacio Moreu, Gustavo M Martins, Ana I Neto, Eva Cacabelos

**Affiliations:** cE3c - Centre for Ecology, Evolution and Environmental Changes/ Azorean Biodiversity Group, and University of Azores, Faculty of Sciences and Technology, Department of Biology, 9501-801 Ponta Delgada, São Miguel, Azores, Portugal; CIBIO - Research Centre in Biodiversity and Genetic Resources, InBIO Associate Laboratory, Pólo dos Açores – Departamento de Biologia da Universidade dos Açores, 9501-801 Ponta Delgada, Portugal; MARE - Marine and Environmental Sciences Centre, Agência Regional para o Desenvolvimento da Investigação Tecnologia e Inovação (ARDITI), Edifício Madeira Tecnopolo, Piso 0, Caminho da Penteada, 9020-105 Funchal, Madeira, Portugal

**Keywords:** macroalgae, oceanic islands, marine bioinvasions, non-indigenous species, Azores

## Abstract

The present study reports the recent occurrence and expansion of *Rugulopteryx okamurae* in the Azores archipelago (NE Atlantic). Morphological and molecular characters confirmed the species identification. Quick surveys around the island of São Miguel showed that it has successfully colonized the island and is quickly expanding. In some locations, *R. okamurae* is currently the dominant organism smothering all other benthic biota and posing a serious threat to the benthic ecosystems across the region. The species first record dates from 2019 near the main harbour of the island, suggesting that its introduction was driven by human-assisted transport, via boat ballast waters or adhered to ship hulls and likely originating from the Mediterranean populations that have been proliferating in recent years across the Strait of Gibraltar.

---

The brown seaweed *Rugulopteryx okamurae* (E.Y. Dawson) I.K. Hwang, W.J. Lee & H.S. Kim 2009, which has been recently introduced in the Mediterranean (Verlaque et al. 2009) is a native and widely distributed species in the subtropical to temperate western Pacific Ocean (Lee and Kang 1986; Silva et al. 1987; Yoshida 1998). It grows in the shallow subtidal and can be present year-round as dormant rhizoidal bases. At macroscopic level, individuals of *R. okamurae* are characterized by a membranous and erect dichotomously branched thallus with smooth margins of 10-20 cm high, attached by rhizoids restricted to the basal parts of the thallus. The fronds grow into a compressed flabellum and in dense groups, and intraspecific variation is known to occur (Hwang et al. 2009). Microscopically, the thallus consists of a medulla, one cell thick centrally and two to three cells thick near the margins (Womersley 1987; De Clerck et al. 2006; Hwang et al. 2009). The reproductive structures are evenly distributed over the entire thallus surface, rather than being confined to the concavities as in the other *Rugulopteryx* species. Just like *Rugulopteryx* species, sporangia of *R. okamurae* are borne on two stalk cells. It can readily form propagules, proliferous branchlets arising on the thallus surface which grow into new plants of the same ploidy level. Although it may be confused with species of the genus *Dictyota*, such as *Dictyota pinnatifida, D. dichotoma, D. spiralis, D. cyanoloma* or *D. fasciola, R. okamurae* is a single taxonomic entity and has been described in Hwang et al. (2009).

In European waters, the non-indigenous *R. okamurae* was first detected in spring 2002 close to a small harbour in Thau Lagoon (France), a known hot spot of marine species introductions in Europe (Verlaque et al. 2009). It was more recently detected in Ceuta and Andalusian waters, southern Iberian Peninsula, in 2015 and 2016, respectively (Altamirano et al. 2016, 2017; Ocaña et al. 2016). In the Strait of Gibraltar, *R. okamurae* has expanded massively in recent years across subtidal marine hard-bottoms, causing evident ecological impacts on coastal communities (García-Gómez et al. 2018, 2020). For instance, after its first record in the area, *R. okamurae* became the most abundant species in only one year (>90% coverage at 10-20 m depth), followed by a significant change in the structure of benthic communities (García-Gómez et al. 2020; Sempere-Valverde et al. 2021). Additionally, hundreds of tons of *R. okamurae* biomass have been reported to accumulate at coastal bedrocks and/or beaches across the region with serious implications for tourism and public health (Ocaña et al. 2016, García-Gómez et al. 2018, 2020). Fishermen have also reported continuous hooks on fishing nets and a significant reduction in fish captures (Sempere-Valverde et al. 2019).

The present study reports the recent occurrence of *Rugulopteryx okamurae* in the Azores archipelago (NE Atlantic). The species was first observed in early 2019 on the south coast of São Miguel Island and identified has a hitherto unknown species of Dictyotales. A search for the species was conducted throughout the island in 2020 (June-July) and again 2021 (May). Sampling was performed at several locations by snorkelling and/or scuba diving at about 5m depth. At each location, the DAFOR scale [D-dominant (>75%); A-abundant (50-75%); F-frequent (25-50%); 0 - occasional (10-25%); R-rare (<10%)] was applied to measure the relative abundance of the algae species along 10 min transect surveys (Fig. 1). In both field campaigns, live plants were collected, taken to the laboratory for examination, and housed as pressed and liquid specimens at the AZB Herbarium Ruy Telles Palhinha of the University of the Azores.

**Figure 1.**
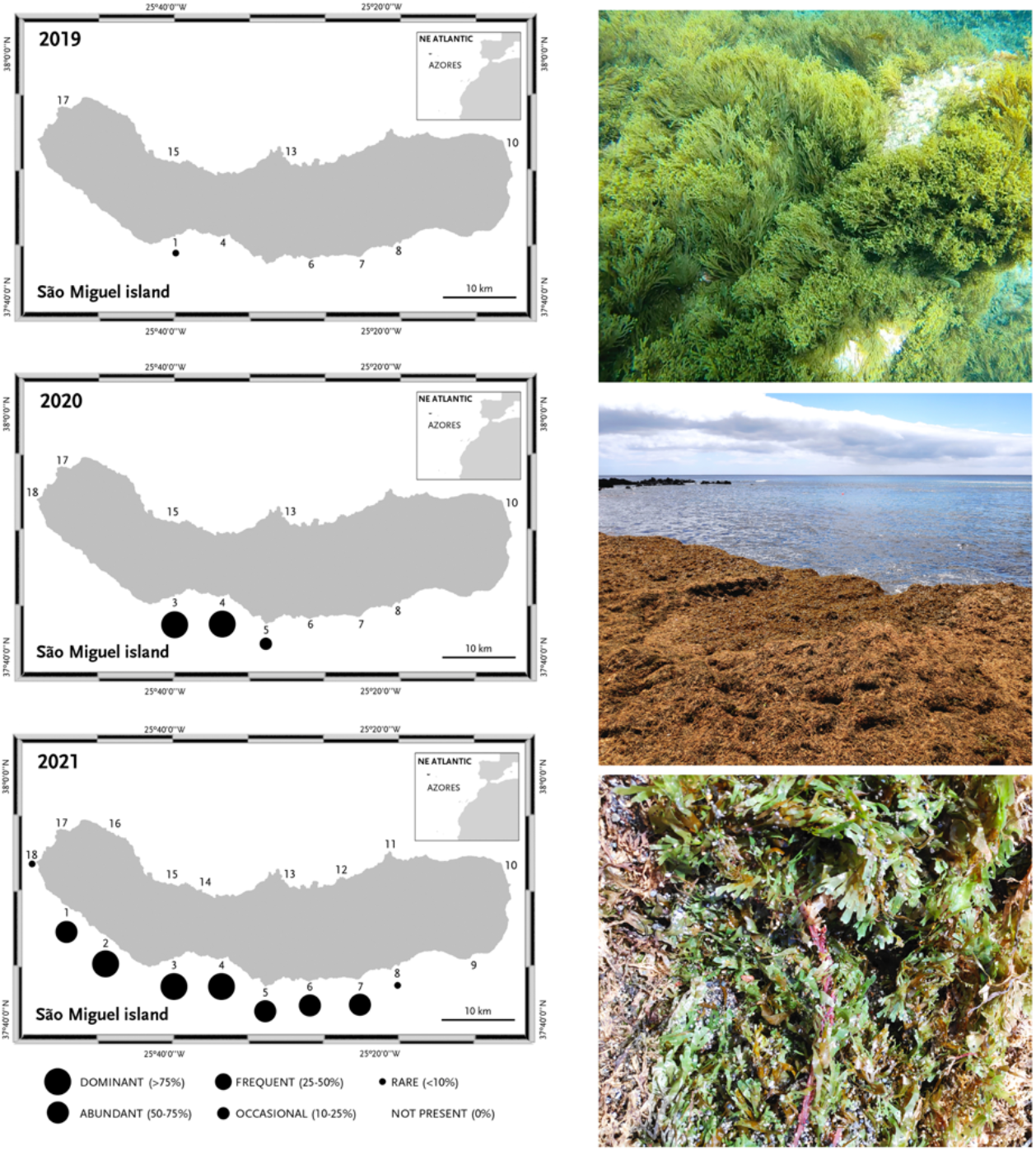
Map of São Miguel Island and locations (numbers) sampled for the presence of *Rugulopterix okamurae* from 2019 to 2021. Black circles indicate the occurrence and relative abundance of the species (see Supplementary material Table S3). Images to the right show *R. okamurae* in its natural environment and as wrack cast away onto the beach.

Collected specimens were identified by their morphological and anatomical characteristics using both binocular and light microscopes (see Hwang et al. 2009; Verlaque et al. 2009). Macroscopically, many individuals displayed numerous proliferous branchlets on both surfaces. Transverse sections revealed the occurrence of mature sporangia subtended by two stalk cells in most individuals, a key distinctive morphologic feature of the genus *Rugulopterix* (Fig. 2). Gametophytes were not encountered in the field.

**Figure 2.**
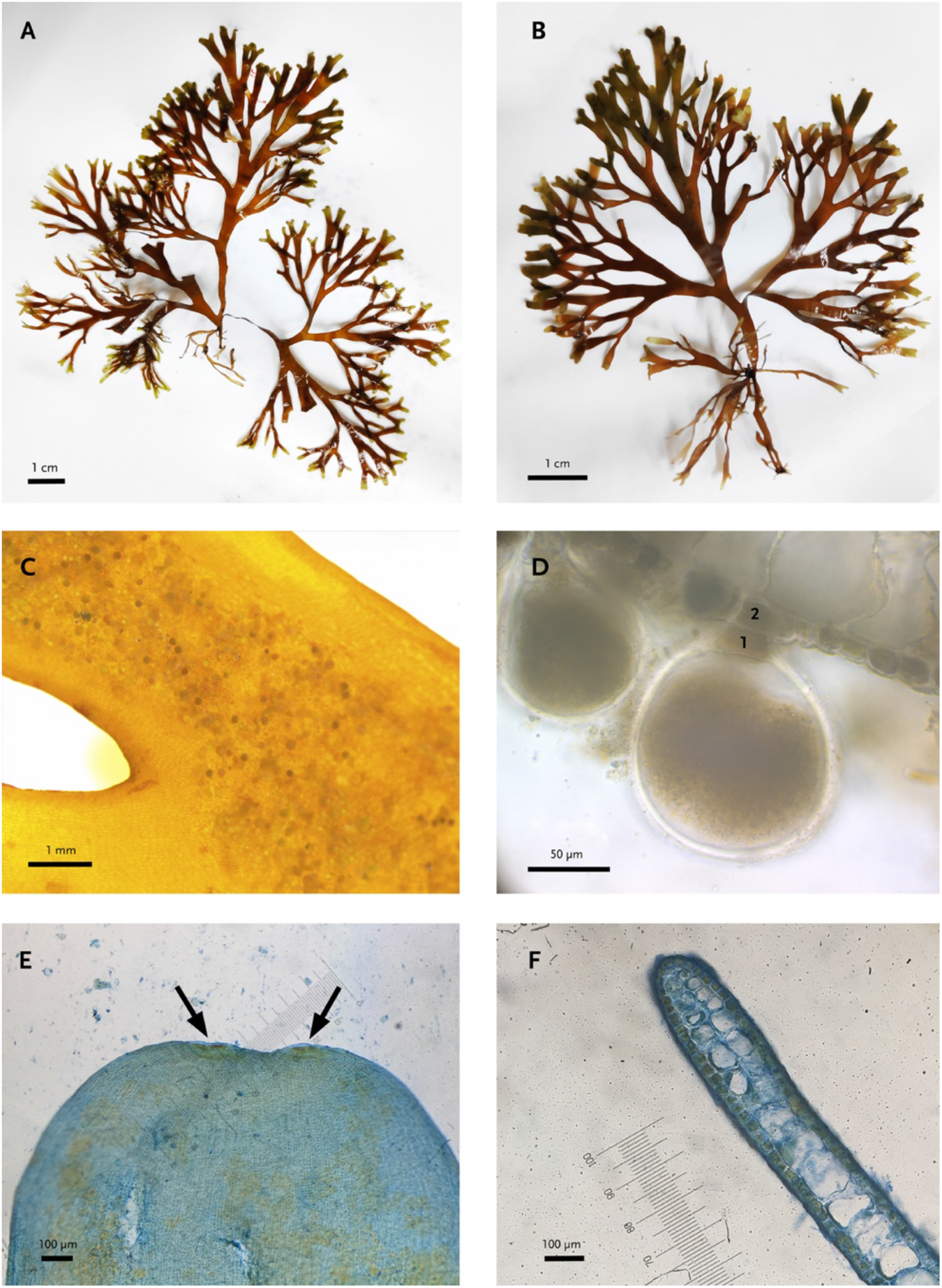
*Rugulopterix okamurae* from the Azores: (A and B) habit of the thallus showing the anchoring rhizoids and dichotomous branching; thallus surface (C) with sporangia (D) that are borne on 2 stalk cells (1 and 2); (E) detail of an apical portion of thallus with two apical cells; a transverse section of thallus showing (F) the margin with two layers of medullary cells. Black bars indicate scale.

Moreover, DNA sequences of two chloroplast protein-encoded genes *rbc*L (ribulose bisphosphate carboxylase large chain) and *psb*A (photosystem II reaction center protein D1) from six collected samples were confirmed as *R. okamurae* using the GenBank BLASTn search. Resulting sequences matched together with *R. okamurae* individuals from western Pacific Ocean (*rbc*L: GenBank accession n° MZ393479 to MZ393484; *psb*A: GenBank accession n° MZ393485 to MZ393490). DNA sequences for *rbc*L and *psb*A were amplified using primers listed in Draisna et al. (2001) and Saunders and Moore (2013), respectively (see Supplementary material for methodological details; Table S1; Table S2).

The arrival of *R. okamurae* in Azores represents its first record in the NE Atlantic islands. Although it may have been initially misidentified as a *Dictyota* spp., the phenotypic characteristics of *R. okamurae* are quite uncommon in the Azores, where most Dyctiotales are smaller in size and exhibit morphological features that can be easily distinguish from *R. okamurae* (e.g. colour, branch dichotomy). A posterior analyses of video data taken in April 2019 indicates that *R. okamurae* may have been already present in the island since early that year. This finding was recorded in a location a few hundred meters from the main shipping port of São Miguel Island, pointing out for the likely vector of introduction of the species in the region. In subsequent years, the species underwent a rapid proliferation throughout the South coast of S. Miguel (Fig. 1, Table S3) dominating subtidal vegetations in both well illuminated and shaded rock surfaces. In some locations occupancy rates were close to 100%. Substantial amounts of the seaweed have also been cast away to the beach as wracks as observed during the first semester of 2021 (Fig. 2). Similar beach accumulations of this brown alga were reported at Faial Island in May 2021, but species identification requires further confirmation.

As mentioned before, and given the location where the species was first detected around the area of the main port of São Miguel, the introduction of *R. okamurae* into the Azores is likely to have been driven by human-assisted transport, via ballast waters and/or fouling (inlays in boat hulls). These are known to be two of the main introduction vectors of exotic species in the marine environment (Hewitt et al. 2009). Both routes could explain the introduction of *R. okamurae* in the region from Mediterranean populations that are proliferating across the Strait of Gibraltar, while taking advantage of the archipelago location as a known stop for transatlantic crossing. For instance, Rosas-Guerrero et al. (2018) observed that adult specimens of *R. okamurae* had survival rates between 80-100% after being grown in dark conditions (a key condition during transport in ballast waters) for three weeks, maintaining the same survival rates when they passed to lighting conditions. Also, it has been shown that the species is capable of adhering to surfaces of very diverse nature and composition (García-Gómez et al. 2018) increasing its capacity to travel attached to boat hulls.

In São Miguel Island, *R. okamurae* is showing explosive growth and a surprising adaptation capacity. Its proliferation and high seabed occupancy rate in São Miguel may be explained by its great ability to propagate by vegetative (i.e. propagules) and asexual mechanisms (i.e. proliferous branchlets arising on the thallus surface which grow into new plants; Verlaque et al. 2009), as highlighted in the Strait of Gibraltar (Altamirano-Jeschke et al. 2017; Altamirano et al. 2019; García-Gómez et al. 2020). Also, anthropogenic disturbances (e.g fisheries, ocean warming) and previous biological invasions (e.g. *Asparagopsis armata* Harvey 1855) (Martins et al. 2019) may have reduced niche competition in the region and favoured the arrival and spread of *R. okamurae* in such remote oceanic islands.

The species is known to cause serious impacts over previously established benthic communities in the Strait of Gibraltar (García-Gómez et al. 2020, 2021; Sempere-Valverde et al. 2021). *R. okamurae* not only has the potential to affect the ecological balance of the community, but also fundamental economic activities such as fishing and tourism. It can contribute to the loss of marine biodiversity and alteration of the structure of the communities, causing the physical displacement of native species due to substrate occupation and preventing the fixation of larvae or propagules of other species. This, together with the amounts of biomass observed at invaded locations, pose a significant ecological pressure to native ecosystems and local economies. This new invader should be kept monitored so that its status and potential impact on Azorean native biota can be evaluated. Given the species high level of proliferation there is a need of a rapid scientific response in providing valuable information so that appropriate preventive measures and mitigation actions can be applied by regional authorities to safeguard coastal marine habitats in Azores.

## Acknowledgments

Funding was provided from National Funds through FCT—Fundação para a Ciência e Tecnologia, within the projects UID/BIA/00329/2015–2019 and UID/BIA/00329/2020–2023. ACLP was supported by PhD grant awarded by FRCT-Fundo Regional da Ciência e Tecnologia (M3.1.a/F/083/2015). In loving memory of Ana Neto.

## Supplementary Material

### Molecular protocol

Specimens used in molecular analyses are listed in Table S1. DNA was extracted using the DNeasy Plant Mini Kit (Qiagen) following the manufacturer’s instructions. Extracted DNA was used as a PCR template consisting of 2 μl DNA, 25 μl MyTaq HS Mix (Bioline), 18 μl sterile H2O, and 0.6 μM of each primer (Table S2) for a total reaction volume of 50 μl. PCR protocols were as follows: i) *rbc*L: 96 °C for 3 min, 27 cycles of 96 °C for 1 min, 42 °C for 2 min, 72 °C for 2 min, followed by a final extension step of 72 °C for 10 min; ii)*psb*A: 94 °C for 2 min, 5 cycles of 94 °C for 30 s, 45 °C for 30 sec, 72 °C for 1 min, 35 cycles of 94 °C for 30 s, 46.2 °C for 30 sec, 72 °C for 1 min followed by a final extension step of 72 °C for 7 min. Amplification products were electrophoresed on a 1.5% agarose gel and positive PCR amplicons were sent to STABVIDA for purification and sequencing. Resulting sequences were edited to eliminate ambiguities and aligned with Bioedit v7.2 (Hall 1999).

**Table S1.** List of specimens collected in São Miguel island used for molecular analyses. Vouchers are deposited in the AZB Herbarium Ruy Telles Palhinha at the University of the Azores.

| Sample code | Location | Date |
| --- | --- | --- |
| SMG-20-07 | Caloura | 27-07-2020 |
| SMG-20-11 | Lagoa | 28-07-2020 |
| SMG-20-13 | Baixa d'Ouro | 29-07-2020 |
| SMG-20-14 | Caloura | 27-07-2020 |
| SMG-21-04 | S. Roque | 17-03-2021 |
| SMG-21-24 | Pópulo | 17-05-2021 |

**Table S2.**
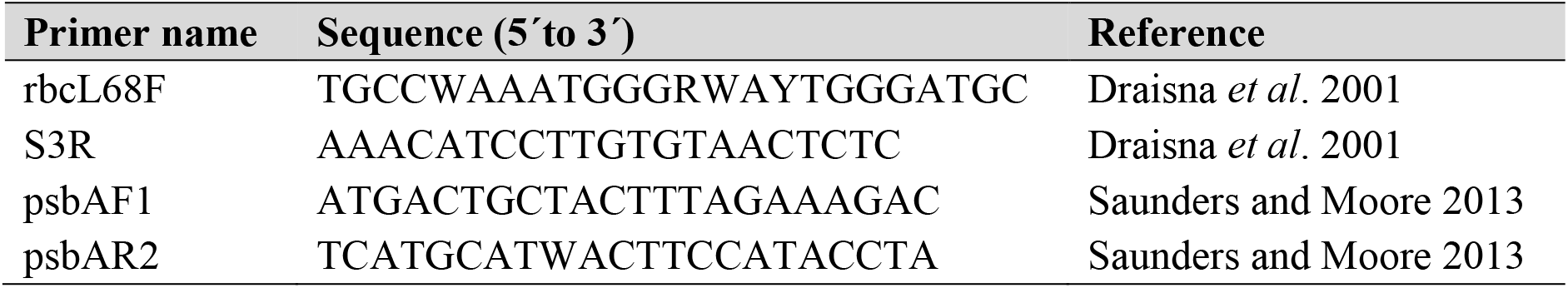
List of oligonucleotides used in this study.

**Table S3.** Field locations surveyed at São Miguel Island in 2019-2021. At each location, the DAFOR scale [D-dominant (>75%); A-abundant (50-75%); F-frequent (25-50%); 0 - occasional (10-25%); R-rare (<10%)] was applied to measure the relative abundance of the algae species along 10 min transect surveys.

| Code | Location | Coordinates (decimal) | 2019 | 2020 | 2021 |
| --- | --- | --- | --- | --- | --- |
| 1 | Feteiras | 37.803488, -25.802394 | Not present | Not present | Abundant |
| 2 | Relva | 37.768793, -25.747248 | Not present | Not present | Dominant |
| 3 | São Roque | 37.743929, -25.639247 | Rare | Dominant | Dominant |
| 4 | Lagoa | 37.741149, -25.573575 | Not present | Dominant | Dominant |
| 5 | Caloura | 37.707311, -25.506511 | Not present | Occasional | Abundant |
| 6 | Vila Franca do Campo | 37.713330, -25.437622 | Not present | Not present | Abundant |
| 7 | Ponta Garça | 37.713824, -25.373018 | Not present | Not present | Abundant |
| 8 | Ribeira Quente | 37.727905, -25.314917 | Not present | Not present | Rare |
| 9 | Faial da Terra | 37.739026, -25.196121 | Not present | Not present | Not present |
| 10 | Nordeste | 37.845703, -25.146526 | Not present | Not present | Not present |
| 11 | Fenais da Ajuda | 37.861622, -25.314817 | Not present | Not present | Not present |
| 12 | Maia | 37.835565, -25.393074 | Not present | Not present | Not present |
| 13 | Ribeirinha | 37.835735, -25.483861 | Not present | Not present | Not present |
| 14 | Calhetas | 37.824471, -25.606583 | Not present | Not present | Not present |
| 15 | São Vicente | 37.834168, -25.668119 | Not present | Not present | Not present |
| 16 | Bretanha | 37.899115, -25.750651 | Not present | Not present | Not present |
| 17 | Mosteiros | 37.899121, -25.814042 | Not present | Not present | Not present |
| 18 | Ferraria | 37.858455, -25.852884 | Not present | Not present | Rare |

## Notes

### Competing Interest Statement

The authors have declared no competing interest.

